# An *Anaplasma phagocytophilum* T4SS effector, AteA, is essential for tick infection

**DOI:** 10.1101/2023.02.06.527355

**Authors:** Jason M. Park, Brittany M. Genera, Deirdre Fahy, Kyle T. Swallow, Curtis M. Nelson, Jonathan D. Oliver, Dana K. Shaw, Ulrike G. Munderloh, Kelly A. Brayton

## Abstract

Pathogens must adapt to disparate environments in permissive host species, a feat that is especially pronounced for vector-borne microbes, which transition between vertebrate hosts and arthropod vectors to complete their lifecycles. Most knowledge about arthropod-vectored bacterial pathogens centers on their life in the mammalian host, where disease occurs. However, disease outbreaks are driven by the arthropod vectors. Adapting to the arthropod is critical for obligate intracellular rickettsial pathogens, as they depend on eukaryotic cells for survival. To manipulate the intracellular environment, these bacteria use Type IV Secretion Systems (T4SS) to deliver effectors into the host cell. To date, few rickettsial T4SS translocated effectors have been identified and have only been examined in the context of mammalian infection. We identified an effector from the tick-borne rickettsial pathogen *Anaplasma phagocytophilum*, HGE1_02492, as critical for survival in tick cells and acquisition by ticks *in vivo*. Conversely, HGE1_02492 was dispensable during mammalian cell culture and murine infection. We show HGE1_02492 is translocatable in a T4SS-dependent manner to the host cell cytosol. In eukaryotic cells, the HGE1_02492 localized with cortical actin filaments, which is dependent on multiple sub-domains of the protein. HGE1_02492 is the first arthropod-vector specific T4SS translocated effector identified from a rickettsial pathogen. Moreover, the subcellular target of HGE1_02492 suggests that *A. phagocytophilum* is manipulating actin to enable arthropod colonization. Based on these findings, we propose the name AteA for *Anaplasma* (*phagocytophilum*) tick effector A. Altogether, we show that *A. phagocytophilum* uses distinct strategies to cycle between mammals and arthropods.

**Importance:** Ticks are the number one vector of pathogens for livestock worldwide and for humans in the US. The biology of tick transmission is an understudied area. Understanding this critical interaction could provide opportunities to affect the course of disease spread. In this study we examined the zoonotic tick-borne agent *Anaplasma phagocytophilum* and identified a secreted protein, AteA, that is expressed in a tick-specific manner. These secreted proteins, termed effectors, are the first proteins to interact with the host environment. AteA is essential for survival in ticks and appears to interact with cortical actin. Most effector proteins are studied in the context of the mammalian host; however, understanding how this unique set of proteins affect tick transmission is critical to developing interventions.

## INTRODUCTION

Multi-host pathogens often have specific adaptation mechanisms to survive in disparate environments (1). For example, vector-borne microbes must adapt and survive in both vertebrate hosts and arthropod vectors (2). These two environments differ significantly with discrepancies in body temperature, nutrient availability, cell types infected, physiological architecture, and immunological pressures (2). Most of our understanding of tick-borne pathogens focuses on interactions with mammalian hosts, as this is where disease occurs. However, the mammal represents only half of the lifecycle for tick borne pathogens. In contrast, little is known about how tick-vectored pathogens mediate interactions with the arthropod. With over 680 million years of evolution separating ticks from mammals (3), our understanding of mammal-pathogen interactions cannot simply be transposed onto ticks (2).

Adapting to different environments is especially critical for obligate intracellular rickettsial pathogens, which are intimately dependent on both arthropod and vertebrate host cells. One of the most common tick-borne rickettsial human pathogens in the United States is *Anaplasma phagocytophilum*, which causes human granulocytic anaplasmosis (4). To complete its lifecycle, *A. phagocytophilum* must transit between *Ixodes scapularis* ticks and mammalian hosts. Interestingly, *A. phagocytophilum* host cell tropisms are not equivalent between mammals and ticks. In mammals, the bacterium preferentially infects neutrophils. In contrast, *A. phagocytophilum* must infect and traverse the tick midgut, travel through the hemocoel, and infect the salivary glands of the arthropod (5, 6). Given the disparities in the host environment and cell tropisms, it is expected that *A. phagocytophilum* would have unique expression profiles adapted for each situation. Transcriptional studies demonstrated that *A. phagocytophilum* differentially transcribes 41% of its genes when growth in tick cells was compared to mammalian cells (7, 8). Transposon mutagenesis found many *A. phagocytophilum* genes are dispensable for growth in the human monocyte cell line HL60 (9), and several of these same genes are upregulated during tick infection. Altogether, this suggests that the tick-specific genes may be involved in arthropod-pathogen interactions (7, 8).

Rickettsial pathogens primarily manipulate host cell biology through effector proteins that are delivered to the host cytosol with a <u>T</u>ype <u>4 S</u>ecretion <u>S</u>ystem (T4SS) (10, 11). T4SS effector molecules subvert host cell defenses and modulate a wide variety of host cell processes (12– 19). A common target of such effectors can be the actin cytoskeleton, which forms the structural scaffolding of the host cell (20). When compared to the model intracellular bacterium *Legionella pneumophila* (1), relatively little is known about the effector repertoire from *A. phagocytophilum* and other rickettsial pathogens. Only five *A. phagocytophilum* T4SS translocated proteins have been identified to date, and none in the context of tick colonization (16, 17, 21–23). The machine learning algorithm OPT4e predicts that *A. phagocytophilum* encodes 48 candidate effectors (24). Fifteen of these candidate genes show unique expression patterns during mammalian and tick cell infection (7, 8). Putative effector HGE1_02492 (APH_0546 in *A. phagocytophilum* strain HZ) demonstrated the highest transcriptional increase when grown in tick cell culture, relative to mammalian cells. Herein, we show that HGE1_02492 is critical for growth in tick cells and colonization of ticks *in vivo*. Further, HGE1_02492 is deliverable by the T4SS into host cell cytosol, where it associates with cortical actin filaments, through multiple sub-domains of the protein. Based on our findings we propose the name AteA, for *Anaplasma* (*phagocytophilum*) tick effector A, and will henceforth refer to HGE1_02492 as AteA. Altogether, we have identified and characterized the first arthropod-vector specific effector from *A. phagocytophilum*, which targets the eukaryotic cytoskeleton.

## RESULTS

An protein of 1103 amino acids (∼120 kD) is encoded by *ateA* (HGE1_02492) and appears to be unique to *A. phagocytophilum*. It has 100% sequence identity with APH_0546 from a different strain of *A. phagocytophilum*, strain HZ. *In silico* analysis of AteA predicts most of the protein is highly disordered, with an N-terminal globular domain (25) and there are two regions containing tandem repeats starting at amino acids 431 and 702 (26).

### Expression of ateA is specific to growth in tick cells

Previous transcriptomics studies using tiling arrays demonstrated that *ateA* was upregulated during *A. phagocytophilum* culture in ISE6 tick cells, and was minimally expressed during growth in the human monocyte-like cell line HL60 (7, 8). Repetitive sequences, like those found in *ateA*, can artifactually inflate transcriptional signals in tiling arrays. We therefore used qRT-PCR with primers targeting non-repetitive sequences to quantify *ateA* expression patterns from *A. phagocytophilum* grown in HL60 and ISE6 cells. Housekeeping *A. phagocytophilum* genes, *rpoB* and *groEL*, are equally expressed when grown in either mammalian or tick cell lines (7, 27), and were therefore used as baseline controls for expression. Our findings demonstrate that *ateA* expression was >8 fold higher during growth in tick cells than in mammalian cells (Figure 1A).

**Figure 1.**
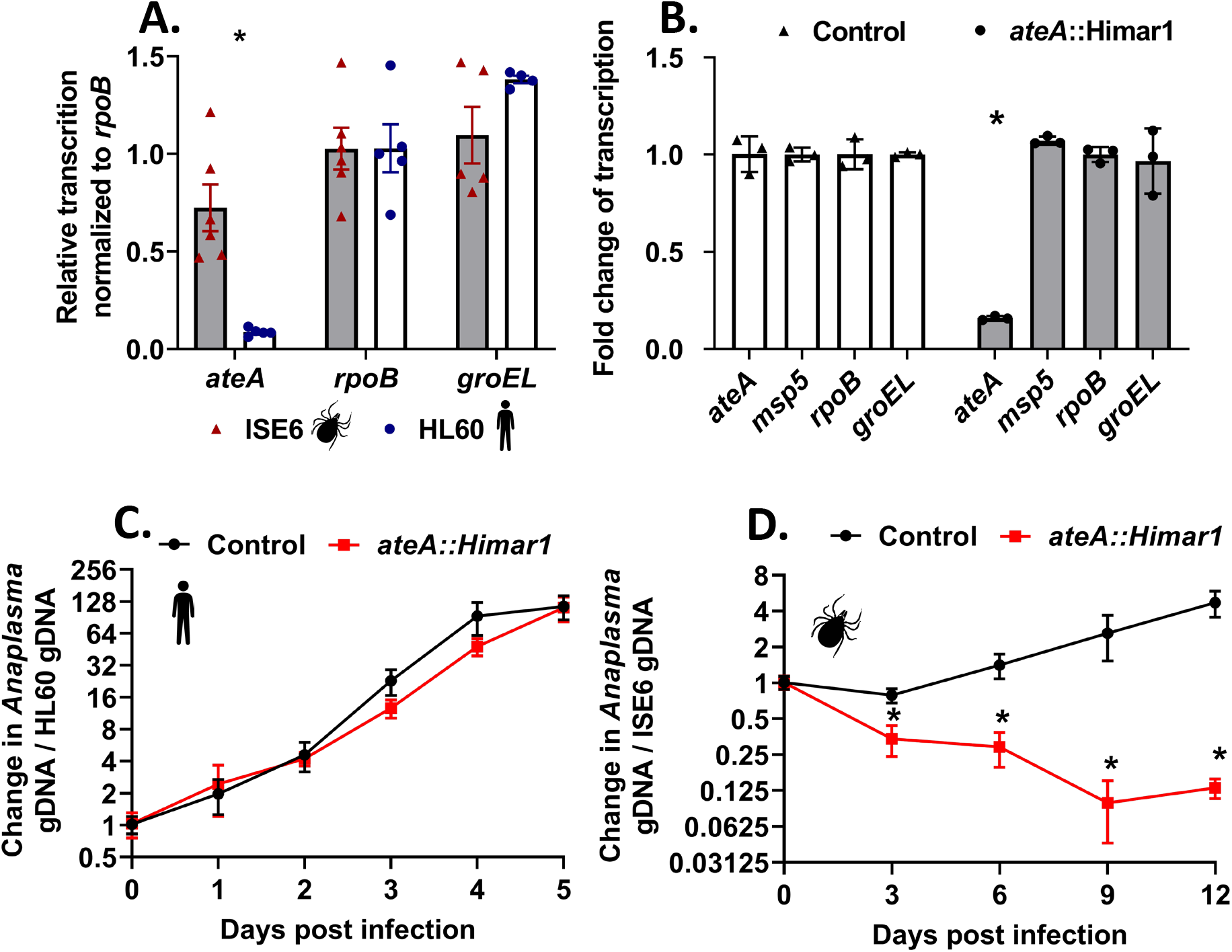
*ateA* is essential for *A. phagocytophilum* survival in tick cells, but dispensable within human cells. A) *A. phagocytophilum* gene expression during growth within tick ISE6 and human HL60 cells. Transcription of *ateA*, and housekeeping genes *rpoB* and *groEL*. B) Transcriptional analysis from *A. phagocytophilum* transposon mutants with the insertion site in a neutral intergenic location (control), or within the *ateA* gene, during culture of with tick ISE6 cells. Transcription measured by qRT-PCR of *ateA, msp5, rpoB*, and *groEL*. Transcription normalized to *rpoB*. Results shown are the mean of three biological replicates with two technical replicates each ± SD. *, P<0.05 (Student T-test). C,D) Growth of *A. phagocytophilum ateA* or control strain in C) human HL60 cells and D) tick ISE6 cells. Bacterial burden was measured as *Anaplasma* gDNA vs host cell gDNA via qPCR. Graphs are representative of three experimental replicates. Data shown are the mean of three biological replicates with two technical replicates each ± SD and is representative of three experimental replicates. *, < 0.05 (Mann Whitney T-test).

### A. phagocytophilum survival in tick cells is dependent on ateA expression

The tick-specific expression pattern of *ateA* led us to ask if it impacts growth in tick cells. For this experiment, we used several tools available to us, including an *ateA* insertion mutant that we previously isolated from a *A. phagocytophilum* Himar1 transposon mutant library (9). As a control strain, we used another mutant, which contains the Himar1 transposon in an intergenic location. This strain has been shown to be phenotypically equivalent to wild-type *A. phagocytophilum* (27, 28). Our analysis confirmed that transcription of *ateA* was abolished in the *ateA*::Himar1 mutant, but transcription of housekeeping genes, *rpoB* and *groEL*, and the conserved *Anaplasma* gene encoding major surface protein 5, *msp5*, were all unaffected (Figure 1B). To test whether the *ateA*::Himar1 mutation impacted growth in the ISE6 cell line, both HL60 and ISE6 tick cells were infected with the *ateA*::Himar1 mutant and growth rates were compared to the intergenic transposon control strain. We found that *ateA*::Himar1 growth in HL60 cells was comparable to the control strain (Figure 1C), but *ateA*::Himar1 growth in ISE6 cells was significantly attenuated, indicating *ateA* is necessary for survival in tick cells (Figure 1D).

### Expression of ateA is necessary for in vivo tick colonization

To examine the importance of *ateA in vivo*, we infected mice with either the control (intergenic transposon strain) or the *ateA*::Himar1 mutant strain. No colonization defect was observed in mice, indicating that *ateA* is dispensable during mammalian infection (Figure 2A). In contrast, larval *I. scapularis* ticks that fed to repletion on *A. phagocytophilum* burden-matched mice acquired significantly less of the *ateA*::Himar1 mutant when compared to the control. This indicates that *ateA* is critical for *A. phagocytophilum* colonization of the tick (Figure 2B).

**Figure 2.**
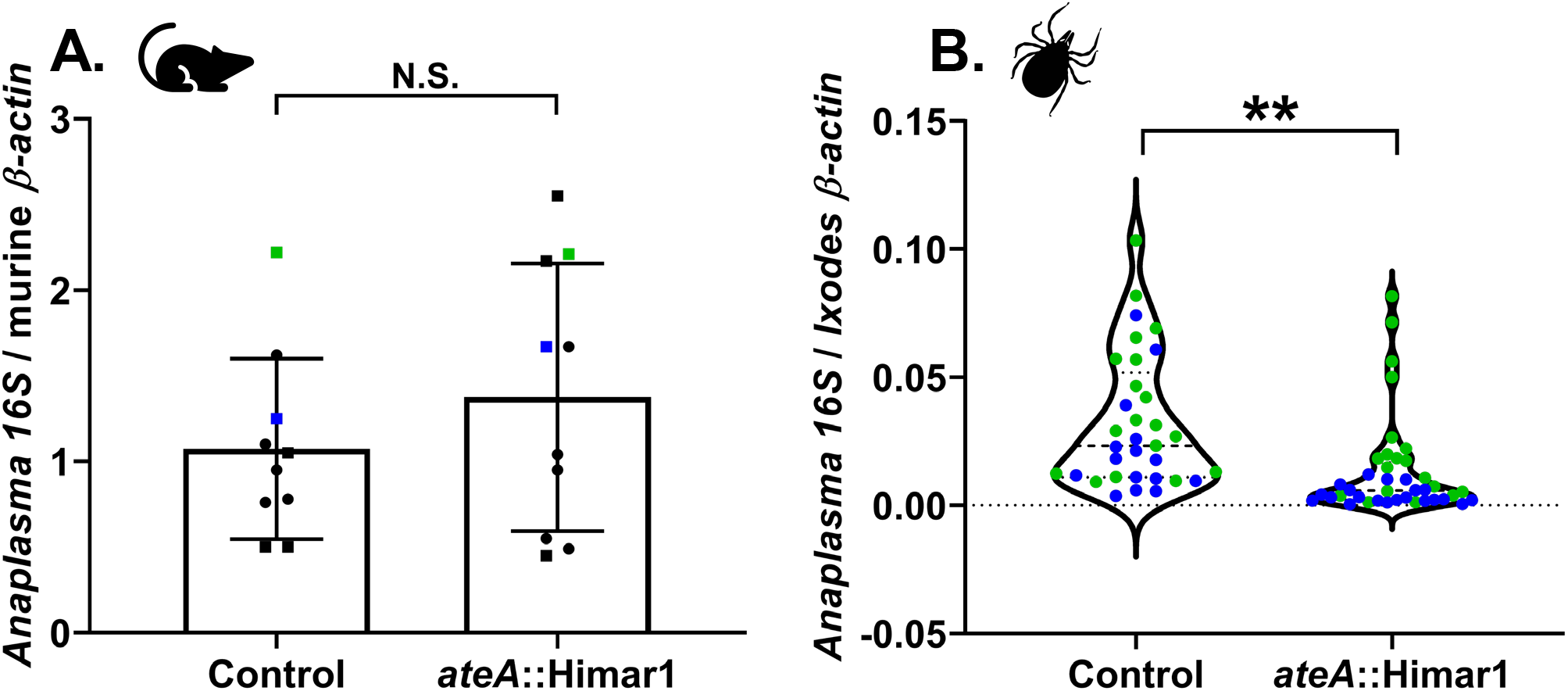
*ateA* is dispensable for murine infection, but mutation attenuates tick acquisition. A) *Anaplasma* burden in mouse blood 7 days post infection by intraperitoneal inoculation with 1×10^8^ host cell free *A. phagocytophilum ateA*::Himar1 or control. Blood samples were processed for DNA isolation and bacterial burden was measured by qPCR of *A. phagocytophilum* 16S rDNA relative to mouse actin by ΔΔCt. Each strain was tested in 5 male (squares) and 5 female (circles) mice and samples were tested in duplicate. Mice used for tick feeding are indicated by blue and green symbols. Blue symbols indicate experimental replicate 1. Green symbols indicate experimental replicate 2. B) *Ixodes scapularis* larvae were infected by feeding to repletion on *A. phagocytophilum* burden-matched mice infected. Whole replete *I. scapularis* larvae were processed for RNA. Bacterial loads were measured by *A. phagocytophilum* 16S rRNA levels relative to mouse actin via qRT-PCR. Data shown represents ticks from two burden matched mouse pairings indicated in blue and green for two experimental replicates. Blue symbols indicate experimental replicate 1. Green symbols indicate experimental replicate 2. 17-20 individual ticks were collected from each mouse as biological replicates, and each qRT-PCR was performed in duplicate. **, < 0.005 (Welsh’s T-test).

### AteA is a T4SS substrate

Several translocated effector prediction algorithms (OPT4e (24), S4TE (29), and T4EffPred (30)) predict that AteA is a T4SS substrate. To empirically test if AteA is translocatable by a T4SS, we used a well-established (31) surrogate assay in *Legionella pneumophila* (32). In this system, the candidate T4SS substrate is fused to adenylate cyclase (CyaA) and expressed in *L. pneumophila*. Candidate effector translocation is detected by accumulation of cAMP in host cells during *L. pneumophila* infection. CyaA-AteA led to significantly greater cAMP than the control (CyaA alone) (Figure 3A). Secretion was not detected from the T4SS deficient *L. pneumophila* strain (*dotA*-), indicating translocation of AteA is T4SS dependent.

**Figure 3.**
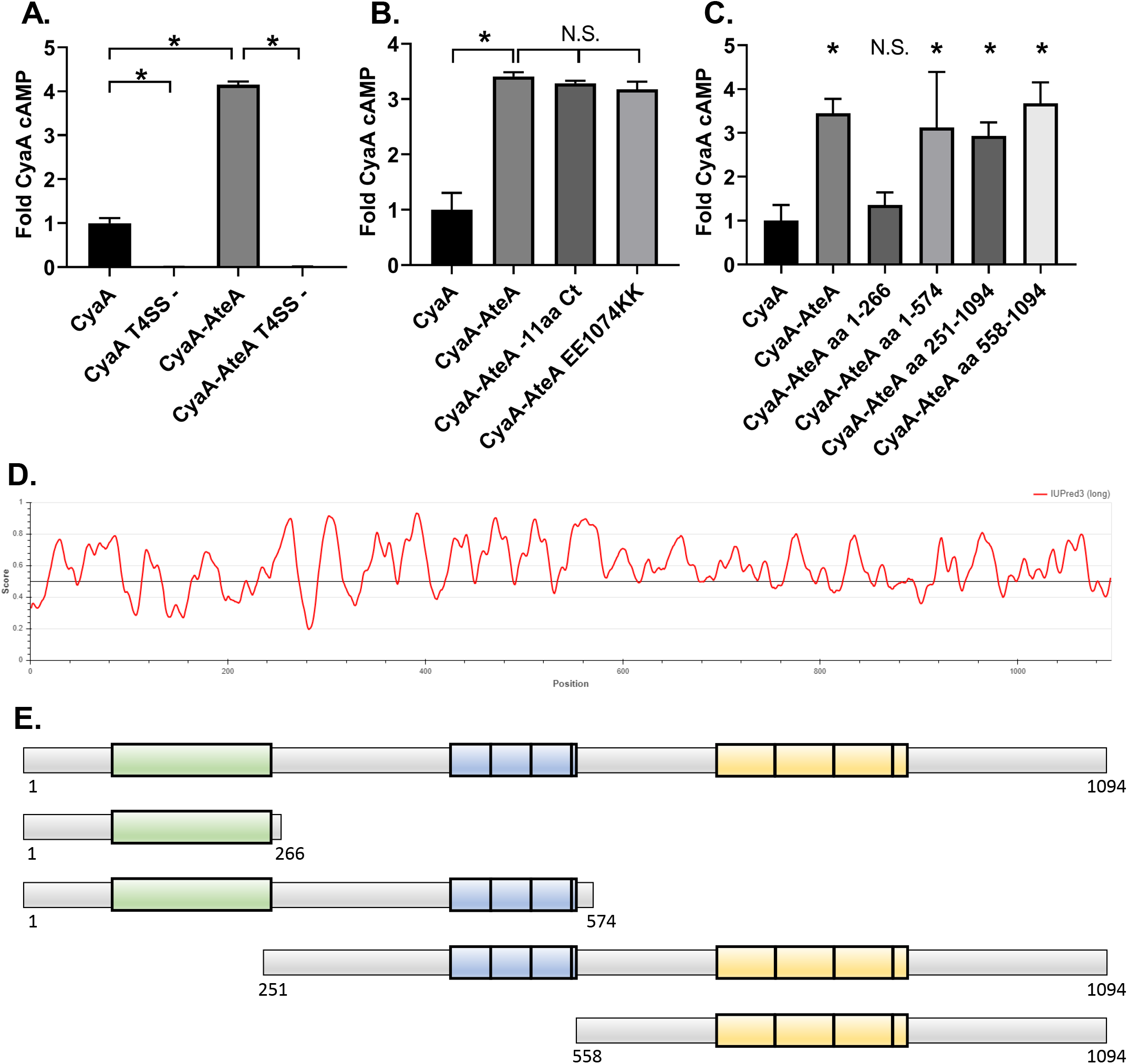
AteA is recognized and secreted by a T4SS. A, B, C) THP-1 cells were infected with a *L. pneumophila* strain expressing the indicated Cya-fusion proteins for 1 hour. cAMP concentrations were quantified from infected cell lysates by ELISA and compared as fold change over CyaA alone. A) CyaA and CyaA-AteA expressed in both wild type *L. pneumophila* Lp02 or T4SS-deficient Lp03 strain (T4SS -). B) C-terminal mutants of CyaA-AteA were expressed in Lp02 and compared to CyaA alone and CyaA-AteA. C) Truncation constructs of CyaA-AteA diagrammed in E were expressed in Lp02 and compared to CyaA alone and CyaA-AteA. D) IUPred3 order/disorder plot of AteA protein. E) Diagram of AteA truncation mutants. Tandem repeat regions highlighted. A, B, C) Error bars represent ± SD of the mean of three biological replicates with two technical replicates each, and graph is representative of two repeated experiments. *, < 0.05 (Mann Whitney T-test).

A secretion signal common to many T4SS translocated proteins are charged residues at the C-terminus (32, 33). We therefore removed 11 C-terminal amino acids from AteA or mutagenized two acidic residues to basic residues in the C-terminus and tested secretion. Neither manipulation strategy affected secretion (Figure 3B), indicating that a different feature of AteA is being recognized by the T4SS. Intrinsically unstructured regions are another feature common among translocated proteins (34). *In silico* analysis of AteA predicts most of the protein is highly disordered, with only the N-terminal portion scoring for a globular structure (25) (Figure 3D). The disordered region has two prominent tandem repeat segments containing either 40 or 59 amino acid repeat units (26) (Figure 3E). We tested four large truncation fragments of CyaA-AteA for translocation (Figure 3E). Truncations retaining a large relative amount of the disordered region were secreted in our assay. The truncation that removed the disordered region, leaving only the N-terminal globular region, was not translocated. Altogether, this suggests that AteA contains multiple internal secretion signals, or that the unstructured nature of the protein itself is being recognized by the T4SS for translocation (Figure 3C,E).

### AteA localizes to the cortical actin cytoskeleton and is dependent on multiple domains

Since AteA is translocated to the host cell, we examined the eukaryotic host cell structures that AteA may be targeting by ectopically expressing a GFP fusion protein (eGFP-AteA) in HeLa cells. Laser-scanning confocal microscopy revealed that eGFP-AteA appeared as branched filamentous structures, resembling the actin cytoskeleton (Figure 4). Filamentous actin (F-actin) was visualized using fluorescently labeled Phalloidin, which revealed that AteA co-localized with actin filaments (Figure 4). Many pathogens are known to target actin, which alters host cell processes with the goal of promoting replication and survival. Two prominent F-actin morphologies in cells are cortical actin and stress fibers. Cortical actin resembles a branched web of fibers just under the cell surface. Stress fibers appear as linear actin bundles connecting two anchor points across the cell (35). The highly branched appearance of AteA localization led us to ask if it was associating with cortical actin. We therefore stained with Cortactin (cortical actin binding protein) and found co-localized with eGFP-AteA (Figure 5). The lack of AteA localization with longer linear actin fibers and the co-localization with Cortactin led us to conclude AteA preferentially associates with the cortical actin network.

**Figure 4.**
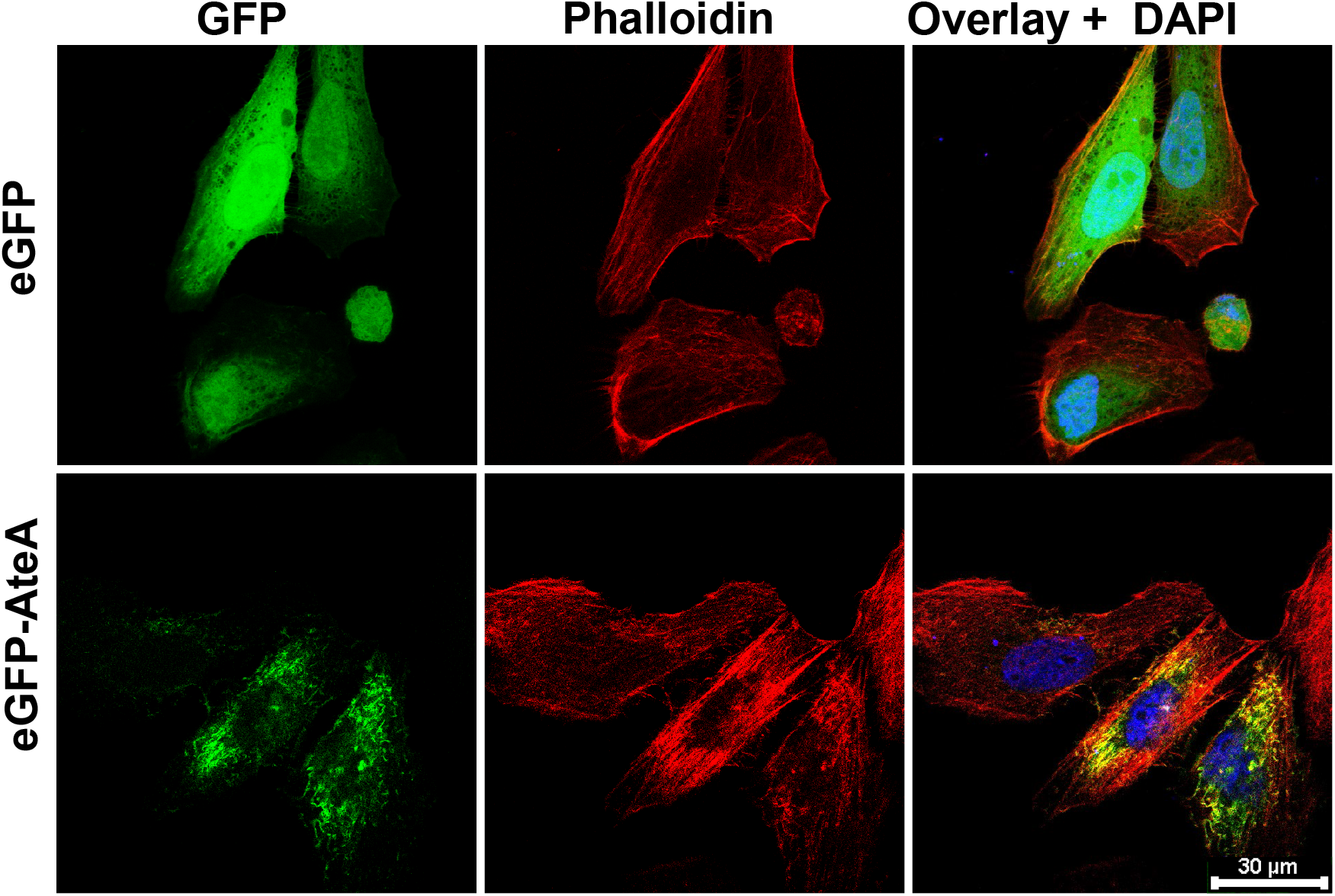
AteA localizes with F-actin. Confocal images of HeLa cells transiently transfected to express eGFP or eGFP-AteA. Actin was stained with Alexa-fluor™ 564 Phalloidin 36 hrs post transfection.

**Figure 5.**
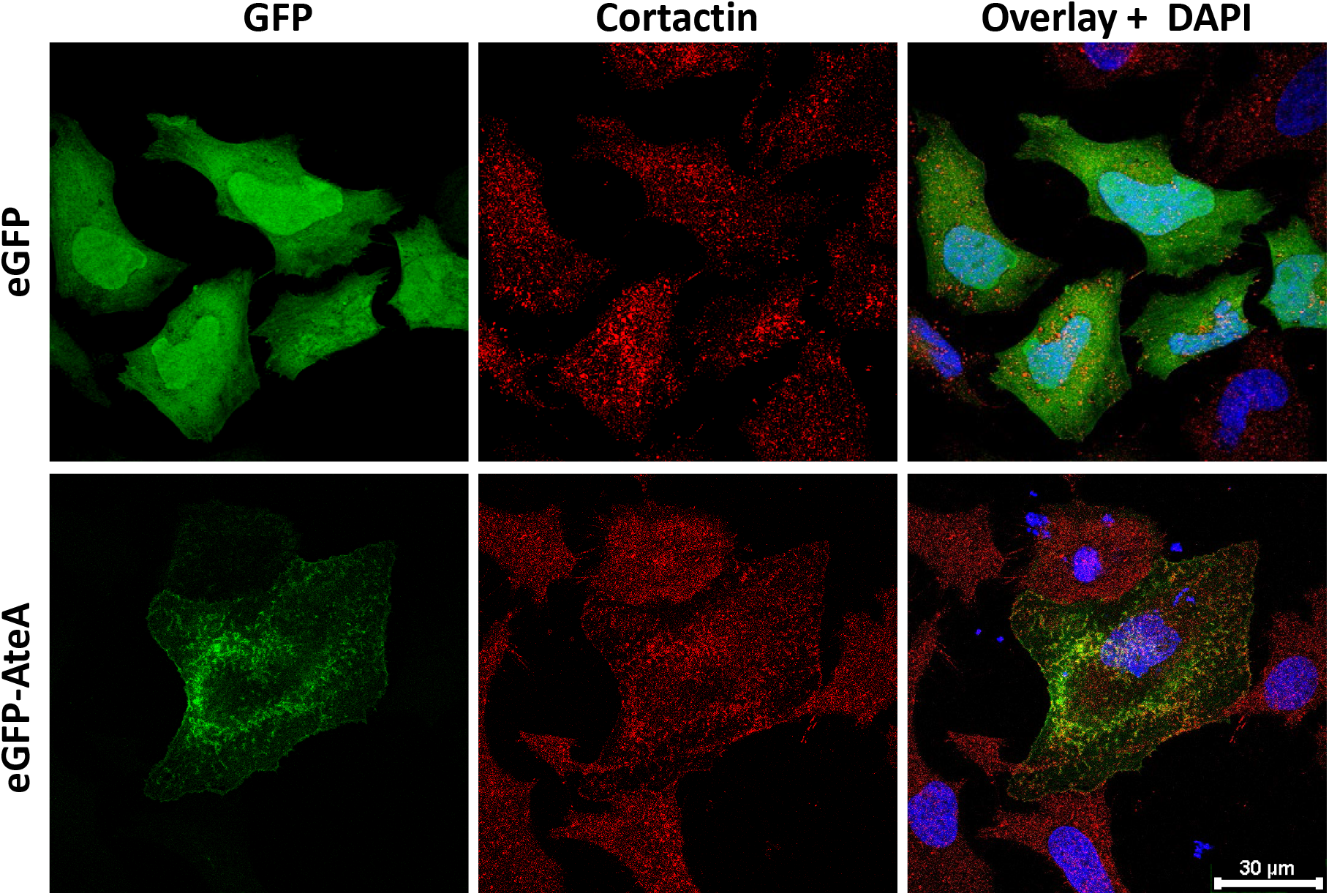
AteA localizes to cortical actin. Confocal images of HeLa cells transiently transfected to express eGFP or eGFP-AteA. Cells were stained to visualize Cortactin at 36 hrs post transfection.

To identify the portions of AteA responsible for localization to the actin cortex, truncations of the protein were constructed and ectopically expressed in HeLa cells (Figure 6). Removal of the C-terminal region following the repeat segments (eGFP-AteA aa1-918) did not change the localization pattern relative to the full-length protein (Figure 6B, C). Truncations that removed the second tandem repeat region (eGFP-AteA aa 1-574) reduced localization with Phalloidin and caused dispersed distribution following the topology of the cell surface (Figure 6D). The N-terminal globular domain alone (eGFP-AteA aa 1-266) showed a similar localization pattern to eGFP-AteA aa 1-574 (Figure 6E). These results suggested that the second tandem repeat region (residues 703-918) is necessary for localization with actin. When this region is removed the protein appears to associate with the cell’s cortex. To test how the other regions of AteA impact the localization we performed the converse experiment, expressing truncations beginning at the N-terminus. AteA lacking the globular N-terminal region (eGFP-AteA aa 251-2094) lost preference for the cell’s cortex, but retained association with actin fibers. This indicates that the N-terminal region of AteA is necessary for the cortical localization (Figure 6F). Interestingly, the actin fibers associated with AteA lacking the N-terminal globular region did not resemble a cortical actin morphology, but appeared more branch-like or distorted than typical actin stress fibers (Figure 6F). Truncations that removed the central region of AteA containing the first set of tandem repeats (eGFP-AteA aa 558-1094) resulted in the loss of the branched pattern altogether. Instead, the protein localized with long linear actin fibers characteristic of stress fibers (Figure 6G). The C-terminal portion of AteA alone (eGFP-AteA aa 918-1094) did not localize with Phalloidin and resembled eGPF alone (Figure 6H). Taken together our findings indicate that the second tandem repeat region of AteA is responsible for localization to actin fibers, the central region of the protein alters this actin localization pattern, and the N-terminal domain provides specificity to the cortex. Altogether, these regions function in combination to associate AteA with actin at the cell cortex.

**Figure 6.**
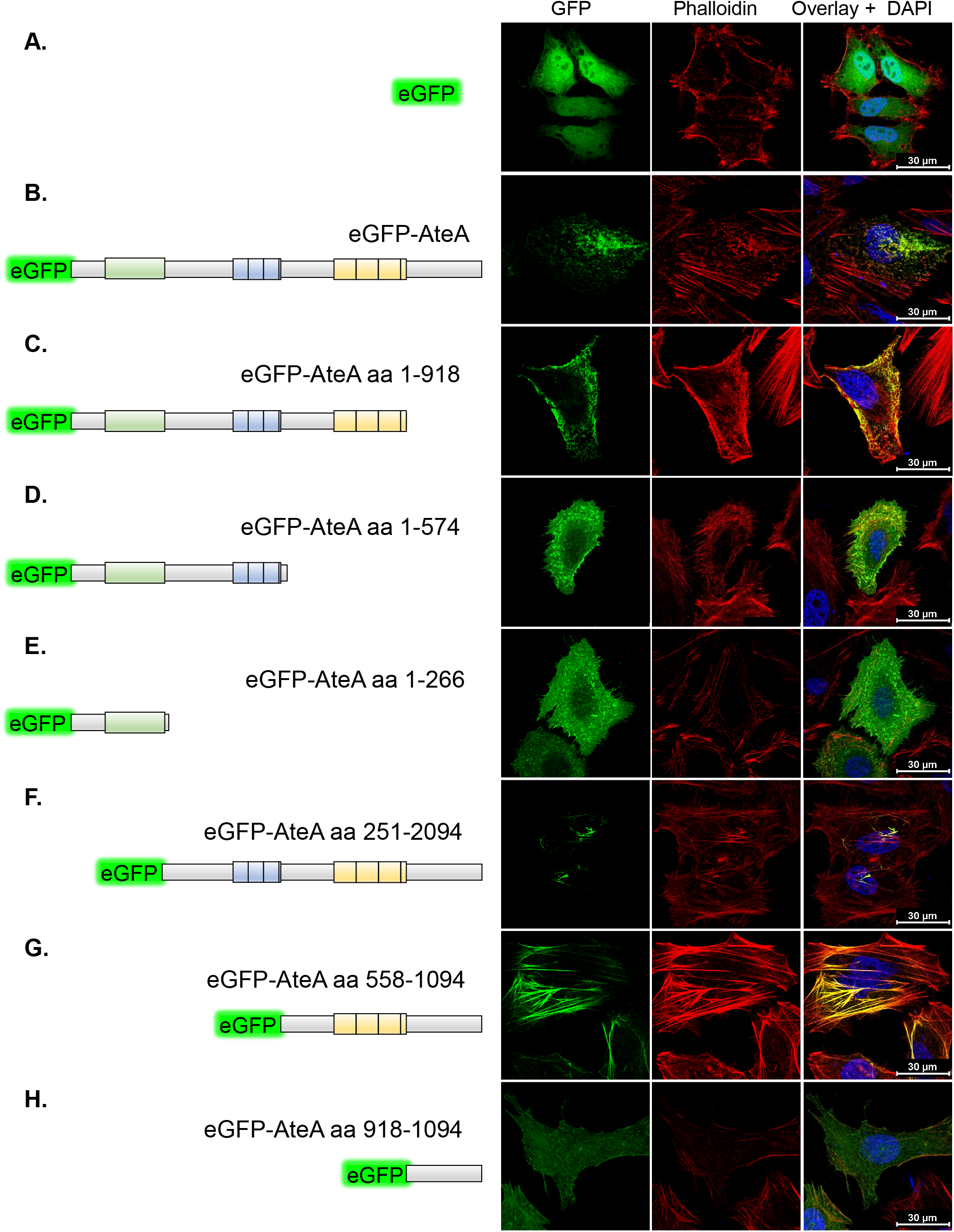
Multiple regions of AteA influence localization with cortical actin. A - H) Left: Schematic of eGFP-AteA fusion constructs and truncations used in transfections. A - H) Right: Confocal images of HeLa cells transiently transfected to express eGFP, eGFP-AteA, or an eGFP-AteA truncation construct as diagramed (Left). Cells were stained to visualize actin using Alexa-fluor™ 564 Phalloidin at 36-48 hrs post transfection.

## DISCUSSION

Here we demonstrate the *A. phagocytophilum* gene encoding *ateA* is highly expressed in the tick environment, is essential for growth and survival within tick cells, and is a T4SS translocated substrate that targets the eukaryotic cytoskeleton. Further, *ateA* is necessary for *A. phagocytophilum* acquisition by *I. scapularis* larvae when feeding on an infected host. However, *ateA* was dispensable for growth in mammalian cell culture, and mutation did not affect bacterial burden in mice. To our knowledge, this is the first description of an arthropod specific rickettsial T4SS translocated effector.

AteA joins the few T4SS effectors identified from *A. phagocytophilum* (16, 17, 21–23). However, machine learning algorithms predict *A. phagocytophilum* encodes many more that remain to be tested (24). Among the 48 putative effectors predicted by OPT4e, fifteen are differentially transcribed between mammalian and tick cells (7, 8). The three best characterized *A. phagocytophilum* T4SS translocated effectors, Ats-1 (17), AnkA (15, 16), and HGE14 (23), are all downregulated during growth in tick cells (7, 8), suggesting their contributions may be more important during mammalian infection. Our understanding of how *A. phagocytophilum* mediates interactions within mammalian cells is limited, but even less is understood about how the bacteria navigate tick cell biology. A full mechanistic understanding of how rickettsial pathogens facilitate their vector-borne life cycle will require effector identification and characterization in the context of both mammalian hosts and arthropod vectors.

While genetic tools among rickettsial organisms remain limited (36), the *A. phagocytophilum* transposon mutant library (9) allowed us to isolate and test a mutation disrupting *ateA*. Although maintenance of the library in HL60 cell culture precludes mutation of genes essential for mammalian infection, it has equipped us to test the contributions of genes that are important for growth in the tick. This is the second mutant from these libraries shown to have a tick cell specific phenotype. Mutation of an outer-membrane O-methyltransferase similarly led to a tick cell specific infection defect (28). Additionally, transposon mutation of a paralogous T4SS component, *virB6-4*, partially attenuated growth in both tick and mammalian cells demonstrating this mutant collection also retains some utility for investigating incomplete phenotypes in mammalian cell models (27). We took the *ateA*::Himar1 mutant beyond cell culture experiments and demonstrated an *in vivo* phenotype through both murine and tick infections that *ateA* is important for bacterial acquisition by ticks from a blood meal. Our work with *ateA*::Himar1 represents the first *A. phagocytophilum* mutant examined in live ticks.

Due to the difficulties of generating recombinant expression systems in an obligate intracellular bacterium (36), efficient T4SS translocation assays using rickettsial organisms have not yet been developed. However surrogate systems in *Legionella* (31), *Coxiella* (23), and *Escherichia* (17, 37) have been used to identify rickettsial T4SS substrates. We demonstrated that *L. pneumophila* recognizes and translocates AteA into the host cell cytosol in a T4SS dependent manner. Motifs at the C-terminus often serve as translocation signals for both the *Legionella* and rickettsial T4SS, but they are not universally required and alternative signals can be used (32, 33). AteA contains multiple charged residues in the C-terminus that we determined are dispensable for translocation. Instead, the large, disordered region of AteA was sufficient for translocation. This suggests that the T4SS is recognizing the unstructured nature of the protein or unidentified internal secretion signals. Indeed, disordered regions are a common characteristic among bacterial effectors (34). While the specificity of the *Legionella* T4SS translocation assay cannot be directly projected onto the *A. phagocytophilum* T4SS, *A. phagocytophilum* effectors Ats1, AnkA, and HGE14 also contain intrinsically disordered regions suggesting that this may be a common feature recognized for T4SS translocation (25).

Intracellular bacteria exist among the scaffold of the host cell’s cytoskeleton composed of actin, tubulin, and various intermediate filaments. Pathogens manipulate this cytoskeleton network to promote internalization, evade destruction, alter intracellular trafficking, and disseminate within and between cells (20). Our understanding of how *A. phagocytophilum* interfaces with this cellular scaffold is limited, but differences exist between mammalian and tick cell infection (38, 39). During mammalian infection the *A. phagocytophilum* vacuole protein AptA recruits intermediate filaments to the *Anaplasma* vacuole. However, transcription of *aptA* is not detectable during *A. phagocytophilum* infection in ticks (39). Actin polymerization is necessary for *A. phagocytophilum* entry into mammalian cells, but during tick cell infection actin is phosphorylated and depolymerized, which is not seen during mammalian infection (38). Although ectopic expression of AteA did not appear to lead to actin depolymerization, this possibility will require further study. We found that AteA localized with branched actin at the cell cortex and was dependent on the predicted tandem repeats and the N-terminal globular domain (Figure 6). Many other intracellular pathogens are known to manipulate cortical actin to attach to the host cell, induce internalization of the bacteria (20, 40), alter endosome maturation (41, 42), block degradation by the lysosome (43), support the pathogen containing vacuole (44), induce extrusion from the host cell (45), and mediate cell to cell bacterial transfer (20, 46). Understanding how AteA manipulates the cytoskeleton to further the *A. phagocytophilum* infection cycle will require further mechanistic dissection.

We demonstrated that *A. phagocytophilum ateA* is specifically important in the tick environment. While examination of AteA localization in tick cells is desired, ectopic expression in ISE6 cells is difficult as transfection efficiency is extremely low, and we were unable to visualize AteA in tick cells. However, we were able to visualize AteA localization with cortical actin in mammalian cells, which may have been possible because actin is one of the most conserved proteins across all eukaryotes (47). Further, mammalian systems have previously been used to investigate multiple actin targeting T4SS effectors, despite mammals not being the evolutionarily relevant environment (1, 41, 42, 48, 49). The tick-specific expression of *ateA* is highlighted by its dispensability during mammalian infection, and severe attenuation of the knockout during tick infection. Why *A. phagocytophilum* requires *ateA* during tick infection remains unclear, but it may stem from the different cell types infected. In mammals the bacteria preferentially infect phagocytic neutrophils, while in ticks it infects multiple non-phagocytic cell types. AteA may be required to induce internalization or trafficking within tick cells, that neutrophils perform without manipulation.

In summary we have identified the first tick-specific translocated effector from *A. phagocytophilum* and have shown that it targets cortical actin. While most research on *A. phagocytophilum* focuses on mammalian infection, it is increasingly clear that the mammalian and tick environments are not equivalent. Given that the arthropod vector is the driver of *A. phagocytophilum* transmission, it is critical to understand how these bacteria survive in the tick. We expect that *A. phagocytophilum* deploys a unique repertoire of effectors to navigate the tick environment, with *ateA* being only the first of many to identify.

## Materials and Methods

### Bacterial and Eukaryotic cell culture

*Escherichia coli* was grown using solid and liquid lysogeny broth (LB) medium with the addition of kanamycin or zeocin 25 μg ml^-1^ antibiotics for selection as needed. *L. pneumophila* Lp02 and Lp03 (dotA−) strains were cultured using N-(2-acetamido)-2-aminoethanesulfonic acid (ACES) buffered yeast extract medium (AYE) and solid charcoal buffered yeast extract agar medium (CYE). *L. pneumophila* cultures were supplemented with 0.4 mg ml^-1^ iron(III) nitrate, 0.135 mg ml^-1^ cysteine (50), 0.1 mg ml^-1^ thymidine, and when appropriate 50 μg ml^-1^ Kanamycin.

HeLa human cervical epithelial cells (American Type Culture Collection [ATCC]; CCL-2) cells were maintained in Eagle’s Minimum Essential Medium (MEM; Corning; 10-010-CV) with 10% fetal bovine serum (FBS: Atlanta biologicals: S11550) and 1× Glutamax (Gibco; 35050061). HL60 human promyelocytic cells (ATCC; CCL-240) and the THP1 human monocyte cell line (ATCC TIB-202) were maintained in Roswell Park Memorial Institute (RPMI) 1640 medium with 10% FBS and 1× Glutamax. Mammalian cell cultures were maintained in a humidified chamber at 37°C with 5% CO_2_. HL60 density was kept between 5 × 10^4^ and 1 × 10^6^ and limited to less than 20 passages to prevent differentiation or phenotypic drift.

*A. phagocytophilum* strain HGE1 and mutant lines were cultured in HL60 cells(9). Insertion mutant *ateA*::Himar1 and control strain were isolated from a previously reported *A. phagocytophilum* Himar1 transposon mutant library (9). The control strain contains the Himar1 transposon in an intergenic location, and has been shown to be phenotypically equivalent to wild-type (27, 28). Infection status of HL60 cells was assessed by Diff-Quick Romanowsky– Giemsa staining. *A. phagocytophilum* were liberated from HL60 cells by 27-gauge needle syringe lysis to generate host-cell-free organisms. Bacterial numbers were estimated as previously described (51, 52).

Tick cells derived from embryonated eggs of the blacklegged tick, *I. scapularis* (Say), ISE6, were grown in L15C-300 medium with 10% FBS (Sigma; F0926), 10% tryptone phosphate broth (TPB; BD; B260300) and 0.1% lipoprotein cholesterol concentrate (MP Biomedicals; 219147680)(53). Infected ISE6 cell cultures were additionally supplemented with 0.25% NaHCO_3_ and 25 mM HEPES buffer (Sigma). Tick cell cultures were incubated at 34°C and 1% CO_2_ (6).

### *Anaplasma phagocytophilum* growth curves

Growth of *A. phagocytophilum* strains in HL60 and ISE6 cells were evaluated similar to previously described (27, 28). Briefly HL60 cells were seeded at 5×10^4^ cells per well of 24 well plates. The plate was then infected with 5×10^4^ host cell-free *A. phagocytophilum* per well for an MOI of 1. Triplicate wells were harvested at the time of inoculation and 1, 2, 3, 4, and 5 days post inoculation. ISE6 cells were seeded at 3×10^5^ cells per well of 24 well plates and allowed to adhere to the plate overnight. Host cell-free preparations of *A. phagocytophilum* strains were prepared immediately before inoculation and bacteria were suspended in L15C300 supplemented with 0.25% NaHCO3 and 25 mM HEPES. ISE6 plates were inoculated at 3×10^6^ *A. phagocytophilum* per well for an MOI of 10. Twenty-four hours post infection the tick cell media was exchanged for fresh L15C300 +NaHCO3 +HEPES to remove remaining extracellular bacteria, and three wells were collected for initial timepoint. Triplicate samples were collected at subsequent time points and frozen to be processed for gDNA using a QIAGEN DNAeasy blood and tissue kit. Change in bacteria and host cell gDNA copies was assessed by qPCR using iTaq universal SYBR green Supermix (Bio-Rad; 1725125) in duplicate reactions. Bacterial gDNA was measured targeting the single copy *A. phagocytophilum* gene *msp5*. HL60 and ISE6 host cell gDNA was measured targeting genes *tlr9* (toll-like receptor 9) and *crt* (calreticulin). Respectively (27) (Table S1).

### Transcriptional analysis of ateA

Triplicate samples were collected during the HL60 and ISE6 *A. phagocytophilum* growth curve experiments. Samples were processed to purify RNA using Direct-zol RNA micro-prep kit^®^ (ZymoResearch) according to product protocols for tissue culture samples. The Verso cDNA Synthesis Kit (ThermoFisher) was used to generate cDNA. Transcripts of *ateA, rpoB, groEL*, and *msp5* genes were measured by qPCR using iTaq universal SYBR green Supermix (Bio-Rad; 1725125) according to Bio-Rad specified cycle conditions. Transcription of *ateA* was compared between experimental groups by ΔΔCt using *rpoB* as the housekeeping control gene.

### Animal infection

Two gender matched groups of ten, 6-week-old C57BL/6 mice (The Jackson Laboratory) were intraperitoneally infected with either the *ateA*::Himar1 mutant or the control strain at 1 × 10^7^ host cell-free *A. phagocytophilum* bacteria per mouse. The control strain contains the Himar1 transposon in an intergenic location, and has been shown to be phenotypically equivalent to wild-type (27, 28). Seven days post infection 25-50 μl of blood was collected from the lateral saphenous vein. Levels of *A. phagocytophilum* in the blood was measured by qPCR (16S rRNA relative to mouse *β-actin* (51, 54)) (Table S1). Uninfected *I. scapularis* larval ticks were purchased from Oklahoma State University (Stillwater, OK, USA). Ticks were housed at 23°C with 16/8-h light/dark photoperiods in 95 - 100% humidity. As sources of tick acquisition for *A. phagocytophilum*, two burden-matched-pairs of mice were selected from the *ateA*::Himar1 and control strain infected mice. Each mouse was individually housed and infested with 200 naïve unfed *I. scapularis* larvae. Three to seven days post infestation replete larvae were collected, individually flash frozen with liquid nitrogen, ground with a pestle, dissolved in TRIzol ^®^ and processed to purify total RNA according to Direct-zol RNA micro-prep kit^®^ protocol. The Verso cDNA Synthesis Kit (ThermoFisher) was used to generate cDNA. Levels of viable *A. phagocytophilum* in the ticks were measured by quantifying *A. phagocytophilum* 16S rRNA relative to *I. scapularis β-actin* transcripts by qRT-PCR (Table S1) by absolute quantification (51, 54). All animal use protocols were approved by the Washington State University Institutional Animal Care and Use Committee (ASAF #6630). The animals were housed and maintained in an AAALAC-accredited facility at Washington State University in Pullman, WA.

### Plasmid Construction

Full length and truncations of the *ateA* open reading frame were amplified from *A. phagocytophilum* genomic DNA with Gateway® compatible primers (Table S1). Amplicons were introduced into pDONR/Zeo by BP Clonase® (Invitrogen). Sequence confirmed inserts were then transferred to destination expression vectors with LR Clonase® (Invitrogen). For ectopic expression in mammalian cells, we used a Gateway® compatible version of pEGFP-C1 (Clontech), pEZYegfp (Addgene). To create a Gateway® destination vector for use in translocation assays (pJC125DEST), the Gateway® attR cassette was inserted into the CyaA translational fusion construct pJC125 (55) at a *SmaI* restriction site. CyaA fusion constructs were introduced to *L. pneumophila* by electroporation.

### Translocation Assay

THP-1 cells were seeded at 5 ×10^5^/mL in 24 well plate and differentiated to macrophage-like cells by treatment with 200 nM Phorbol 12-myristate 13-acetate (PMA; Sigma) for 18 hrs. Transformed *L. pneumophila* cells were grown overnight to an OD_600_ of 2.0, at which point the bacteria were in post-exponential phase, highly motile, and infectious. Expression of the CyaA fusion proteins was induced by adding 1 mM IPTG for 1 hour, and motility was verified by microscopy of wet mounted samples. Cell culture medium was used to dilute *L. pneumophila* which was then used to infect THP-1 cells at an MOI of 1. One hour post infection, cAMP was extracted and quantified as previously described (56) using the cAMP Parameter Assay Kit (R&D Systems).

### Immunofluorescence

HeLa cells were transfected using FuGENE® 6 Transfection Reagent at a 3:1 FuGENE to DNA ratio. Proteins were expressed for 36 – 48 hrs, then fixed in 4% paraformaldehyde. Fixed cells were permeabilized with 0.1% Triton X-100 for 15 min and washed three times in PBS. Cells were incubated with Alexa Fluor™ 568 Phalloidin (Thermo Fisher Scientific) in PBS containing 1% bovine serum albumin (BSA) for 30 min. Cells were washed three times for five minutes with PBS and coverslips were mounted on slides using Vectashield® mounting medium with DAPI. Slides were imaged using a Leica SP8 confocal microscope.

### Statistics

All *in vitro* experiments were performed with three biological replicates, measured in technical duplicate assays, and experiments were repeated three times to ensure reproducibility of findings. *In vivo* experiments used ten independently inoculated mice per group. 17-20 ticks were collected per mouse. Two burden matched mice pairs were used for experimental replicates of the tick feeding. Data were expressed as means and graphed with standard deviation. Data points were analyzed with a Student T-test (Mann-Whitney) for *in vitro* experiments and *in vivo* experiments were analyzed with an unpaired Welch’s T-test. Statistical analysis was performed and graphed with GraphPad Prism version 9.0. A P-value of <0.05 was considered statistically significant.

## ACKNOWLEDGEMENTS

We are grateful to Daniel Voth at the University of Arkansas College of Medicine (Little Rock, AR) and Jean Celli at Washington State University Paul G. Allen School for Global Health (Pullman, WA) for sharing protocols and plasmids constructs for the CyaA secretion assay. Tamara O’Connor at Johns Hopkins University (Baltimore, MD) generously provided *L. pneumophila* Lp02 and Lp03 (*dotA*^−^) strains. Lisa Price and Nicole Burkhardt at the University of Minnesota (Saint Paul, MN) for their assistance in isolating and verifying strains from the Himar1 transposon collection.

## AUTHOR CONTRIBUTIONS

**Jason M. Park:** Conceptualization, Methodology, Investigation, Writing - Original Draft & Editing, Visualization, Funding acquisition. **Brittany M. Genera:** Methodology, Investigation, Visualization, Reviewing & Editing. **Deirdre Fahy:** Methodology, Investigation, Review & Editing. **Kyle T. Swallow:** Investigation. **Curtis M. Nelson and Jonathan D. Oliver:** Investigation, Resources. **Dana K. Shaw:** Methodology, Investigation, Review & Editing. **Ulrike G. Munderloh:** Resources, Funding acquisition. **Kelly A. Brayton:** Conceptualization, Supervision, Funding acquisition, Review & Editing.

**Supplemental Table 1.**
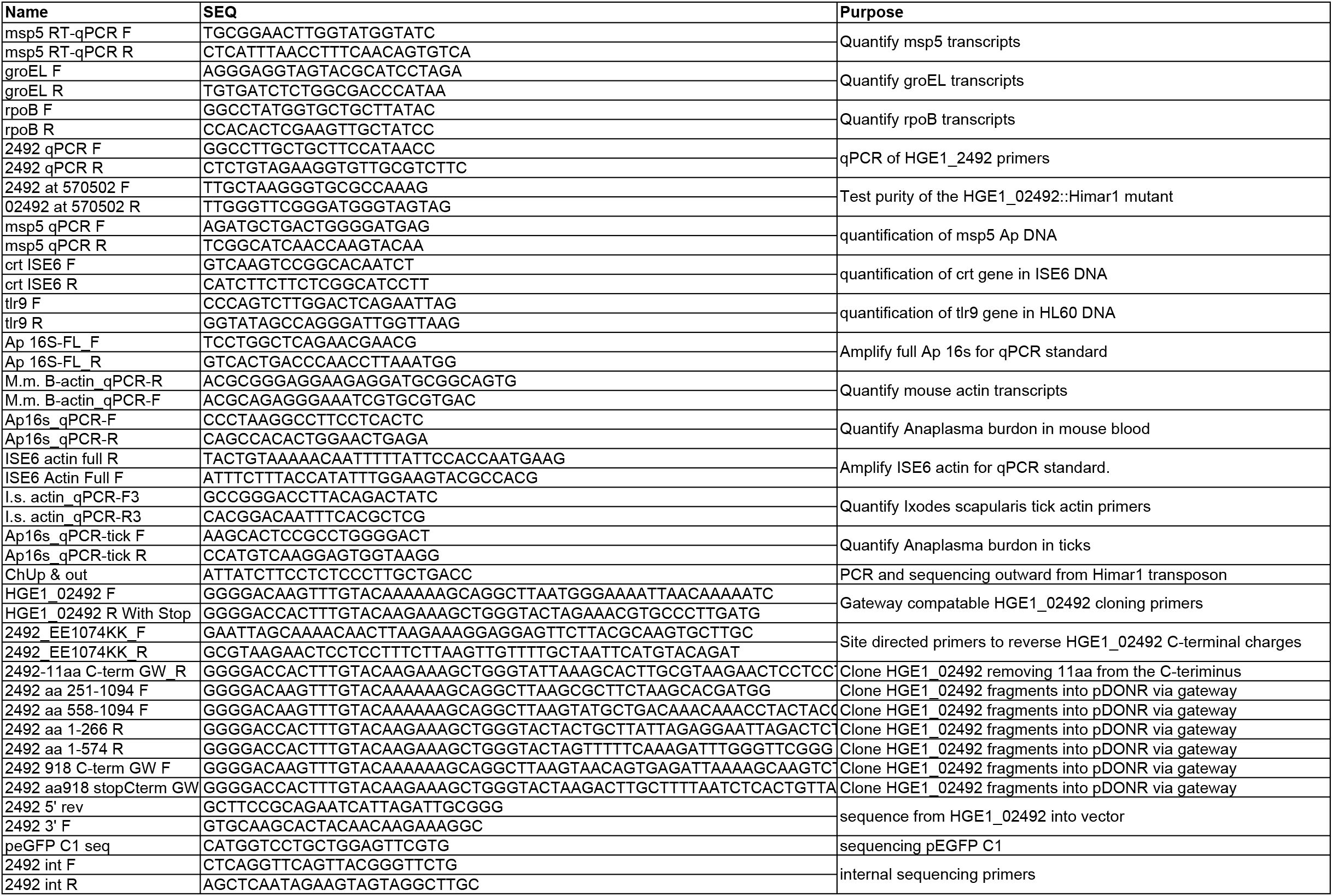

