## Supplemental Table 1 for "An *Anaplasma phagocytophilum* T4SS effector, AteA, is essential for tick infection"

| Name | SEQ | Purpose |
|---|---|---|
| msp5 RT-qPCR F | TGCGGAACTTGGTATGGTATC | Quantify msp5 transcripts |
| msp5 RT-qPCR R | CTCATTTAACCTTTCAACAGTGTCA | |
| groEL F | AGGGAGGTAGTACGCATCCTAGA | Quantify groEL transcripts |
| groEL R | TGTGATCTCTGGCGACCCATAA | |
| rpoB F | GGCCTATGGTGCTGCTTATAC | Quantify rpoB transcripts |
| rpoB R | CCACACTCGAAGTTGCTATCC | |
| 2492 qPCR F | GGCCTTGCTGCTTCCATAACC | qPCR of HGE1_2492 primers |
| 2492 qPCR R | CTCTGTAGAAGGTGTTGCGTCTTC | |
| 2492 at 570502 F | TTGCTAAGGGTGCGCCAAAG | Test purity of the HGE1_02492::Himar1 mutant |
| 02492 at 570502 R | TTGGGTTCGGGATGGGTAGTAG | |
| msp5 qPCR F | AGATGCTGACTGGGGATGAG | quantification of msp5 Ap DNA |
| msp5 qPCR R | TCGGCATCAACCAAGTACAA | |
| crt ISE6 F | GTCAAGTCCGGCACAATCT | quantification of crt gene in ISE6 DNA |
| crt ISE6 R | CATCTTCTTCTCGGCATCCTT | |
| tlr9 F | CCCAGTCTTGGACTCAGAATTAG | quantification of tlr9 gene in HL60 DNA |
| tlr9 R | GGTATAGCCAGGGATTGGTTAAG | |
| Ap 16S-FL_F | TCCTGGCTCAGAACGAACG | Amplify full Ap 16s for qPCR standard |
| Ap 16S-FL_R | GTCACTGACCCAACCTTAAATGG | |
| M.m. B-actin_qPCR-R | ACGCGGGAGGAAGAGGATGCGGCAGTG | Quantify mouse actin transcripts |
| M.m. B-actin_qPCR-F | ACGCAGAGGGAAATCGTGCGTGAC | |
| Ap16s_qPCR-F | CCCTAAGGCCTTCCTCACTC | Quantify Anaplasma burdon in mouse blood |
| Ap16s_qPCR-R | CAGCCACACTGGAACTGAGA | |
| ISE6 actin full R | TACTGTAAAAACAATTTTTATTCCACCAATGAAG | Amplify ISE6 actin for qPCR standard. |
| ISE6 Actin Full F | ATTTCTTTACCATATTTGGAAGTACGCCACG | |
| I.s. actin_qPCR-F3 | GCCGGGACCTTACAGACTATC | Quantify Ixodes scapularis tick actin primers |
| I.s. actin_qPCR-R3 | CACGGACAATTTCACGCTCG | |
| Ap16s_qPCR-tick F | AAGCACTCCGCCTGGGGACT | Quantify Anaplasma burdon in ticks |
| Ap16s_qPCR-tick R | CCATGTCAAGGAGTGGTAAGG | |
| ChUp & out | ATTATCTTCCTCTCCCTTGCTGACC | PCR and sequencing outward from Himar1 transposon |
| HGE1_02492 F | GGGGACAAGTTTGTACAAAAAAGCAGGCTTAATGGGAAAATTAACAAAAATC | Gateway compatable HGE1_02492 cloning primers |
| HGE1_02492 R With Stop | GGGGACCACTTTGTACAAGAAAGCTGGGTACTAGAAACGTGCCCTTGATG | |
| 2492_EE1074KK_F | GAATTAGCAAAACAACTTAAGAAAGGAGGAGTTCTTACGCAAGTGCTTGC | Site directed primers to reverse HGE1_02492 C-terminal charges |
| 2492_EE1074KK_R | GCGTAAGAACTCCTCCTTTCTTAAGTTGTTTTGCTAATTCATGTACAGAT | |
| 2492-11aa C-term GW_R | GGGGACCACTTTGTACAAGAAAGCTGGGTATTAAAGCACTTGCGTAAGAACTCCTCC | Clone HGE1_02492 removing 11aa from the C-teriminus |
| 2492 aa 251-1094 F | GGGGACAAGTTTGTACAAAAAAGCAGGCTTAAGCGCTTCTAAGCACGATGG | Clone HGE1_02492 fragments into pDONR via gateway |
| 2492 aa 558-1094 F | GGGGACAAGTTTGTACAAAAAAGCAGGCTTAAGTATGCTGACAAACAAACCTACTAC | Clone HGE1_02492 fragments into pDONR via gateway |
| 2492 aa 1-266 R | GGGGACCACTTTGTACAAGAAAGCTGGGTACTACTGCTTATTAGAGGAATTAGACTC | Clone HGE1_02492 fragments into pDONR via gateway |
| 2492 aa 1-574 R | GGGGACCACTTTGTACAAGAAAGCTGGGTACTAGTTTTTCAAAGATTTGGGTTCGGG | Clone HGE1_02492 fragments into pDONR via gateway |
| 2492 918 C-term GW F | GGGGACAAGTTTGTACAAAAAAGCAGGCTTAAGTAACAGTGAGATTAAAAGCAAGTC | Clone HGE1_02492 fragments into pDONR via gateway |
| 2492 aa918 stopCterm GW | GGGGACCACTTTGTACAAGAAAGCTGGGTACTAAGACTTGCTTTTAATCTCACTGTTA | Clone HGE1_02492 fragments into pDONR via gateway |
| 2492 5' rev | GCTTCCGCAGAATCATTAGATTGCGGG | sequence from HGE1_02492 into vector |
| 2492 3' F | GTGCAAGCACTACAACAAGAAAGGC | |
| peGFP C1 seq | CATGGTCCTGCTGGAGTTCGTG | sequencing pEGFP C1 |
| 2492 int F | CTCAGGTTCAGTTACGGGTTCTG | internal sequencing primers |
| 2492 int R | AGCTCAATAGAAGTAGTAGGCTTGC | |
